# Transfer learning in heterogeneous drug-target interaction predictions using federated boosting

**DOI:** 10.1101/2023.01.14.524052

**Authors:** Dániel Sándor, Péter Antal

**Affiliations:** Department of Measurement and Information Systems, Budapest University of Technology and Economics, Budapest, Hungary

**Keywords:** federated learning, multitask learning, boosting, DTI

## Abstract

In multitask federated learning, when small amounts of data are available, it can be harder to achieve proper predictive performance, especially if the clients’ tasks are different. However, task heterogeneity is common in modern Drug-Target interaction (DTI) prediction problems. As the data available for DTI tasks are sparse, it can be challenging for clients to synchronize the tasks used for training. In our method, we used boosting to enhance transfer in the multitask scenario and adapted it to a federated environment, allowing clients to train models without having to agree on the output dimensions. Boosting uses adaptive weighting of the data to train an ensemble of predictors. Weighting data boosting can induce the selection of important tasks when shaping a model’s latent representation. This way boosting contributes to the weighting of tasks on a client level and enhances transfer, while traditional federated algorithms can be used on a global level. We evaluate our results extensively on the tyrosine kinase assays of the KIBA data set to get a clear picture of connections between boosting federated learning and transfer learning.

## I. Background

In the past years, drug discovery has become more and more reliant on the use of machine learning [1]. For a long time, drug discovery was mostly conducted with in vitro tests on candidates. But these types of tests are expensive and time-consuming to execute thus, they do not scale well for the large amounts of available drug candidates [2]. Machine learning has the advantage of scalability and relatively low resource costs thus, it can complement traditional tests well. Modern drug discovery problems usually involve a model which can select or narrow down a list of drug candidates which are further tested in laboratories [3].

The motivation for our work is a federated drug-target interaction problem. In this, we try to predict which chemical compounds (drugs) will bind to which biological targets (e.g. proteins) in the human body, to induce a change in the organism [4]. For this task, there is already a large amount of data available [5]–[7]. The historically used compounds of drugs usually have multiple targets to which they are likely to bind hence the only job is to find these targets. The data on targets is available through assays. Assays are procedures to analyze the qualities of a given target. In this case, this means the identification of drugs that bind or do not bind to any given target. A usual DTI task is built up in the following way: multiple assays are available for one target, and assays are only conducted for some compounds (and not all available) to save resources. After the collection of assays, they can be arranged into a bioactivity matrix, which contains one assay in a column and one compound in a row. Because the assays only have available data on interactions where they have been measured and the measurements are only conducted where the researchers saw a chance of interaction the resulting data is sparse and missing-not-at-random (MNAR) type data.

To explain the motivation of federated learning (FL), we have to look at the nature of the data. When working with high-value data, like data on drugs, companies can be reluctant to share it. Although in some cases the demand presents itself to learn from a larger data set and cooperate with other companies or research institutions. In these cases, privacy-preserving federated learning can be a useful tool for performing the training. In federated learning, multiple parties agree to train their models together for better performance, while their data sets remain private and not shared. This can be achieved in a multitude of ways. One of the most prominent examples is Federated Averaging (FedAvg) [8], which uses a shared model architecture and averages the models’ weights every few iterations.

Multitask learning is a natural extension of this setup as it enhances the performance of models on targets when learned jointly [9]. However multitask learning also presents challenges, namely selecting tasks in a way to achieve the best possible performance [10]. To solve this we introduce an adaptive weighting by the use of boosting. This way at the beginning of training no task selection is necessary instead the data will be weighted in every iteration to maximize the performance of models.

Boosting is a highly versatile method, thus it can be used for drug research too. In [11] Svetnik et al. showed that tree-based boosting is especially useful in Quantitative structureactivity relationship (QSAR) problems, where the goal is to predict bioactivity from structural features, much like in DTI. However, in an FL scenario boosting is still a relatively new concept and only a handful of solutions exist. Most of these are from the field of gradient boosting: One of these is SecureBoost [12], which is a framework for gradient boosting by decision trees, however, it only works in vertical FL scenarios, when the data is only distributed in the feature space, not in the sample space, which is a strong assumption. In [13] Li et al. create a similarity-based federation for gradient boosting, where each party makes their data public through hashing and they train models on the hashed features. These approaches are significantly different from AdaBoosting, and in the next chapter, we give a detailed explanation, of why AdaBoost is a good fit for the problem at hand. The adaptation of boosting to multitask can also be done in several ways: Wang et al. [14] created the Online Multitask Boosting (OMB) algorithm, which is the generalization of a transfer learning boosting algorithm. Their algorithm captures task relatedness by calculating the differences in errors of the ensemble on the two tasks. In their algorithm, some tasks are used to train models of the ensemble and every task learns to adapt them by assigning its own weight to the output of the classifier. A different approach is described by Zhang et al. [15] in a technique they call MTBoost. In MTBoost they learn the relationships of tasks in the form of a task covariance matrix. First, they learn an ensemble, for a base hypothesis and generate the output for every task by learning the weights specifically for them. The base hypothesis can be formulated by learning an aggregated ”super-task” based on the task covariances.

In our work, we aim to combine the best of all these approaches in a way that makes it feasible to create a multitask ensemble of neural networks in a federated manner, which leverages the adaptive weighting of samples to boost predictive performance. To do this we create a combination of the FedAvg and the AdaBoost algorithms, using neural networks as base classifiers. We present our results on the KIBA data set, with a realistically constructed federated split for the data.

## II. Method

To understand federated boosting, we start from the same setup as the FedAvg algorithm: Every client has their own data set, with possible overlaps in both the compound- and target spaces. In the beginning, the clients assign a uniform weight to each of their compounds: *ω*_*i*0_ = 1/*N*, *i* = 1, 2,…, *N*

After this, the server initializes the model weights *w_t_* and distributes them to the clients. The clients run a local training sequence on the model, in my experiments, this means one epoch of training the network. Next, the clients send their models back to the server for aggregation (alternatively, like in FedAvg it is enough to send the gradients). The server averages the weights and produces the final model, which will be part of the ensemble.

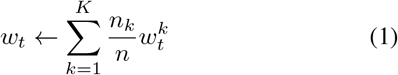

After this, the model is sent back to the clients and they calculate the error of the model on their data using the formula from AdaBoost, with the difference being, that we classify a compound as correctly classified if a given threshold of correctness is reached on its predictions. In my measurements, this meant that 80% of a compound’s measurements are to be predicted correctly for the compound to be classified as correct.

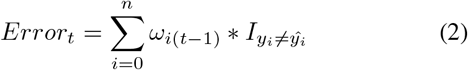

Based on the error, they assign an individual weight to the model (same as AdaBoost).

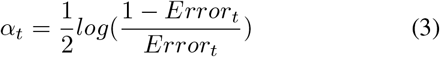

Now the partners know the weight of the model, they can reweight the data and start the training again with new weights from the server.

### Algorithm 1 FedMTBoost

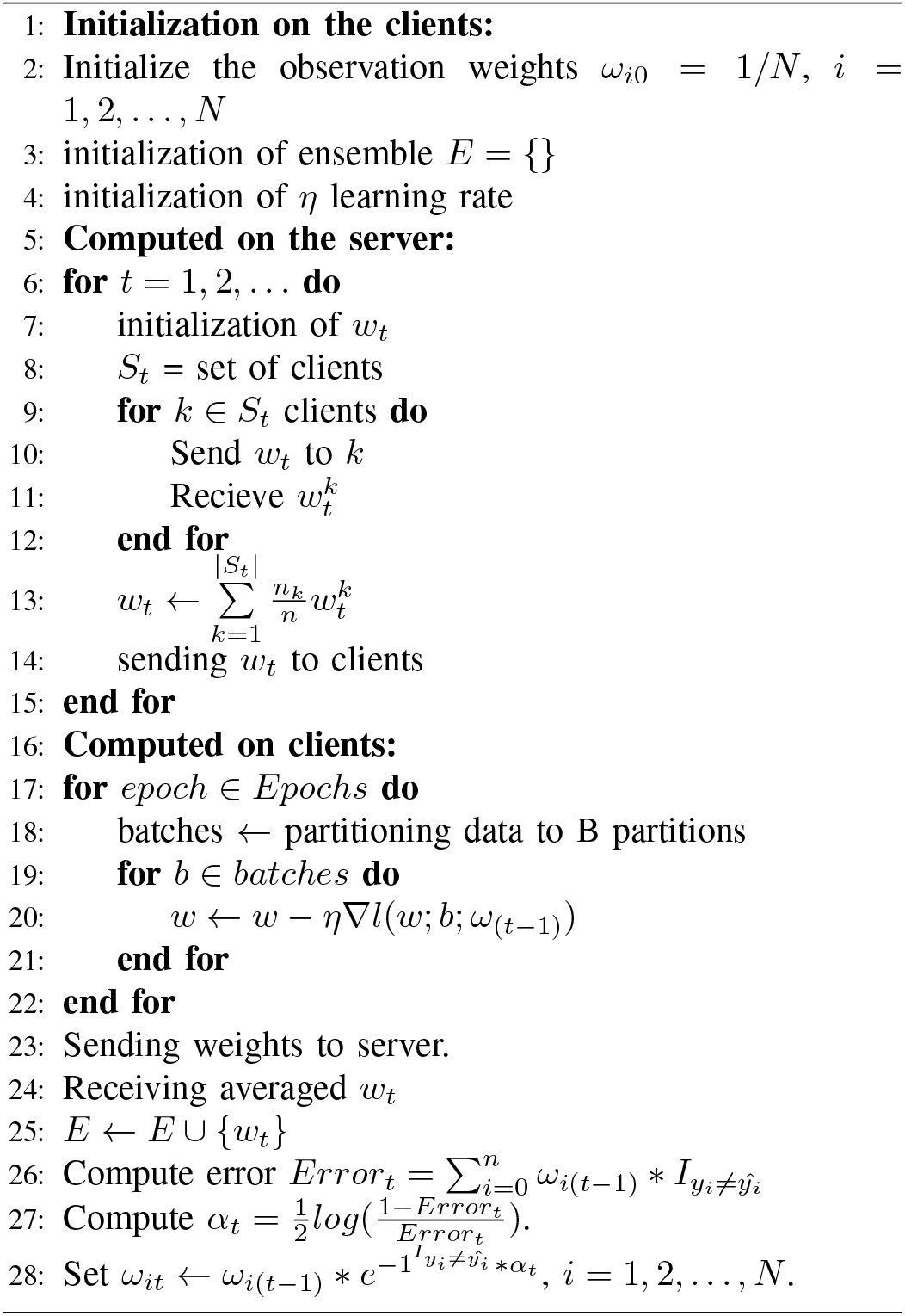

The FedMTBoost algorithm can be viewed from a global point, and this way it is essentially carrying out the same steps as FedAvg, but when it is viewed from the clients’ local point it resembles the AdaBoost algorithm, with the managing of weights during training.

When predicting for new data, every client can use the shared ensemble and create a weighted average using their private model weights.

## III. Results

To make sense of different technologies affecting the results of training and the methods’ dependence on data we have devised a set of experiments to be evaluated on different dataset sizes. The experiments can be split along three axes: singletask and multitask, boosted and non-boosted methods, and finally based on the type of federation: singleparty, multiparty and federated. This resulted in the following experiments (with each having a single- and a multitask variant):

- Singleparty single Multilayer perceptron (MLP) model: All data pooled and one model trained
- Singleparty boosted: All data pooled and AdaBoost ensemble trained
- Multiparty single MLP model: Data distributed to 10 clients and everyone trains an MLP on their part of the data
- FedAvg: Data distributed and partners train MLP in a federated way, with the FedAvg algorithm
- FedMTBoost: Data distributed and clients train an ensemble with the FedMTBoost algorithm

### A. Technical details

The methods were evaluated on the tyrosine kinase assays of the KIBA dataset, which resulted in 60 tasks unevenly distributed in the multiparty scenarios. The MLP was based on the SparseChem [16] implementation with one hidden layer with a size of 1400, using Adam optimizer. The federation of data happened both in the compound and the assay spaces, which means that averaging is only possible in the first layer, and not on the output (as different clients might use entirely different assays). We compare the AUROC scores of the methods on a five-fold dataset, with fold ”0” always being used for evaluation. For folds 1, 2, 3 and 4, every possible subset is used for training, with the complementary folds being used for evaluation too.

### B. Comparing singleparty methods

To see what effect boosting has on a baseline MLP method we first evaluated them on the whole dataset. The results below show the performances of boosting and MLP-based training, which as we can see in this scenario is not significant. The results are similar in a singletask training, but with worse performances.

**TABLE I.**
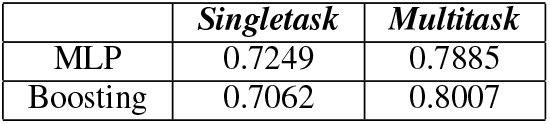
Singleparty average AUROC scores for 4 fold trainings

|  | *Singletask* | *Multitask* |
| --- | --- | --- |
| MLP | 0.7249 | 0.7885 |
| Boosting | 0.7062 | 0.8007 |

### C. Comparing multiparty methods

The more interesting results come, when the data is distributed and not every client has access to every assay and compound. First to get a baseline every client trains a model for themselves without federation, and this can be compared with the federated methods instead of a singleparty performance.

When looking at federated averaging we can see that for partners with enough data it does improve the predictive performance, which is expected as more information is present for these types of models, and they are able to develop a better representation. Although the improvement is present it is not statistically significant in most relevant cases. As a significance threshold of *p* = 0.05 would be required.

The case for FedMTBoost is different: It can improve the predictive performance of FedAvg and multiparty MLPs.

The improvements here are in most cases significant and are present in both multitask and singletask scenarios. In Fig. 4 we can see the performance of partner 8, which is one of the larger partners, but it can demonstrate the usual scale of improvements.

**Fig. 1.**
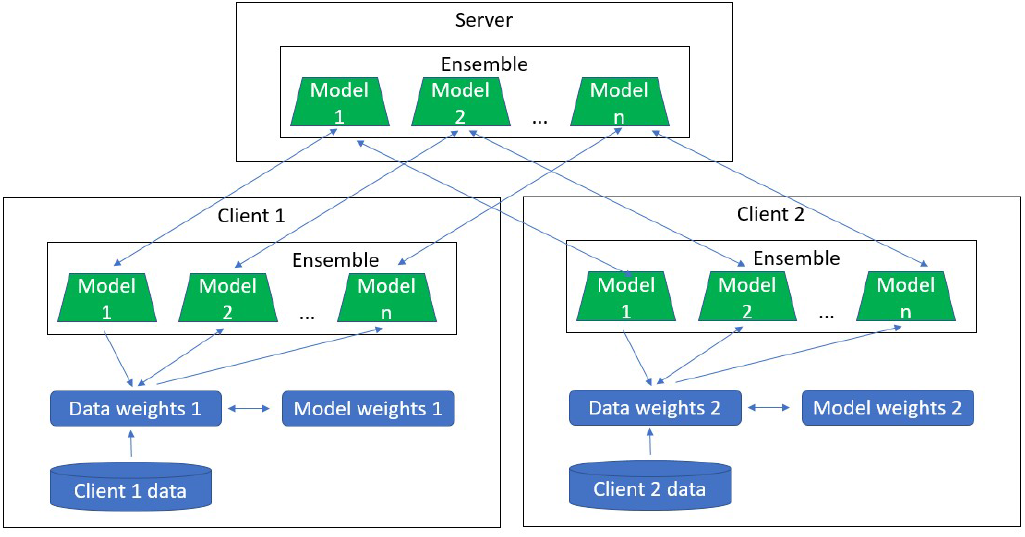
High-level overview of FedMTBoost.

**Fig. 2.**
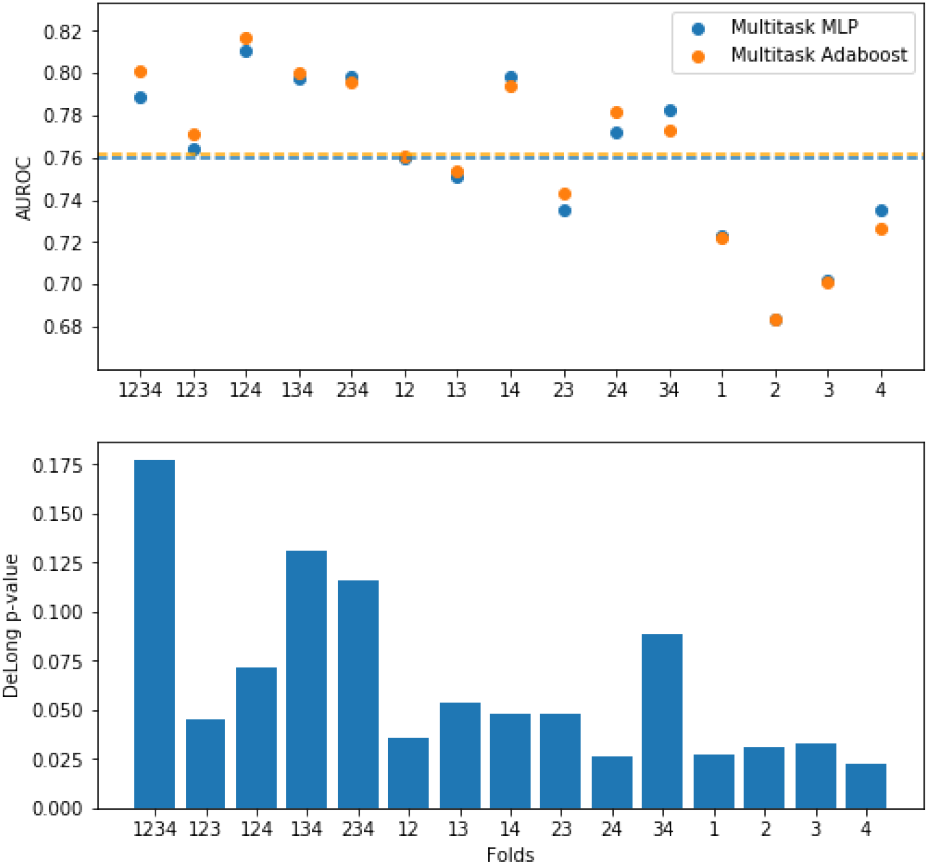
AUROC scores and DeLong p-values of singleparty methods for every set of folds used in training.

**Fig. 3.**
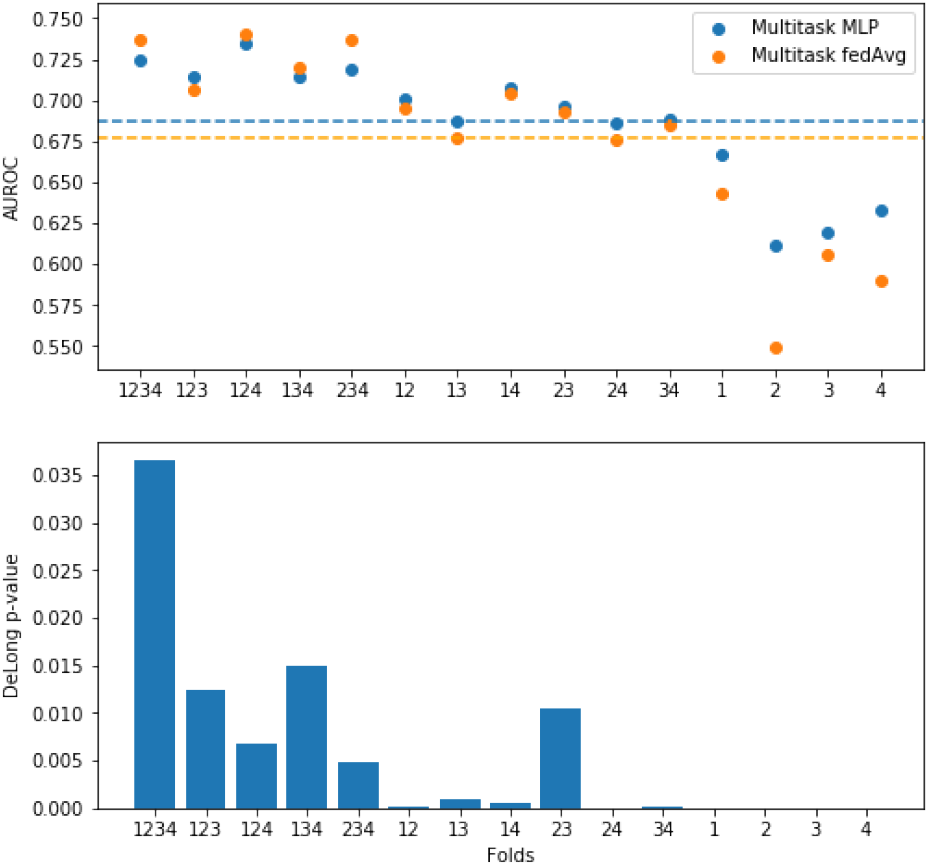
AUROC scores and DeLong p-values of multiparty and federated methods for every set of folds used in training.

**Fig. 4.**
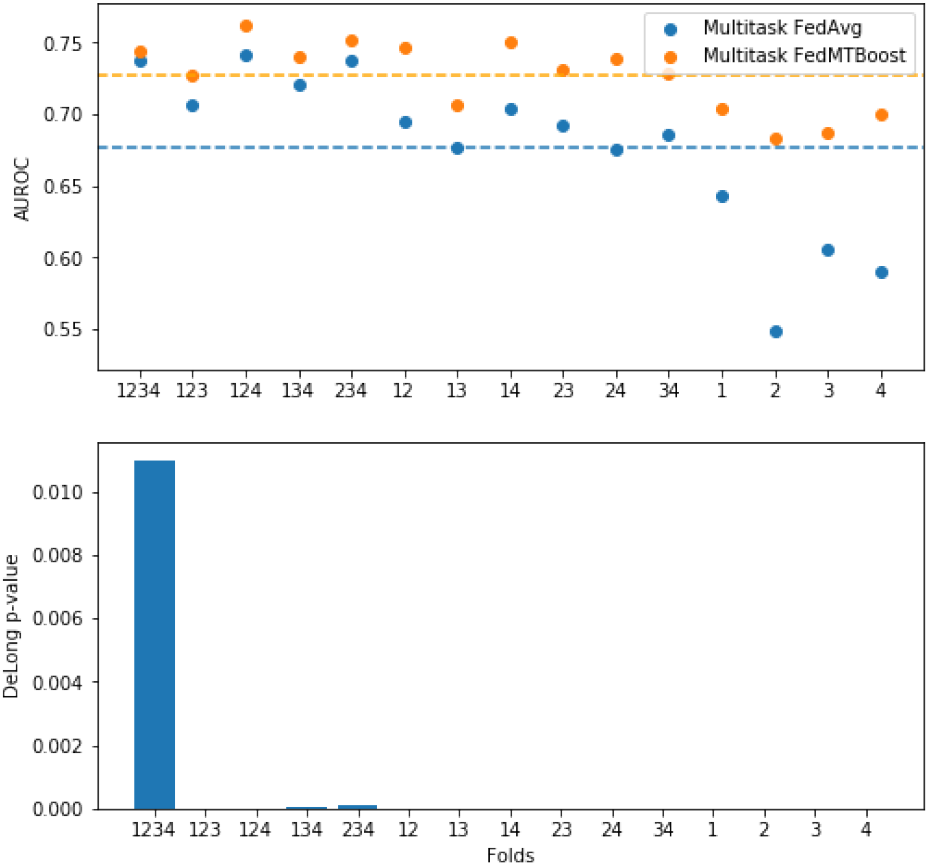
AUROC scores and DeLong p-values of federated methods for every set of folds used in training.

It is important to note that the most significant differences were achieved when not all the data was present for training.

## IV. Conclusion

As we showed Federated Boosting is a possible and useful approach that is adaptable to both single- and multitask environments. It does improve the predictive performance of FedAvg by leveraging data weighting from boosting. This proves that ensemble methods are a good fit to use in federated learning.

**TABLE II.** Multiparty average AUROC scores for 4 fold trainings on partner 8

|  | *Singletask* | *Multitask* |
| --- | --- | --- |
| Multiparty MLP | 0.6625 | 0.725 |
| FedAvg | 0.6235 | 0.7374 |
| FedMTBoost | 0.7325 | 0.7445 |

